# Transient kinetic analysis of SWR1C-catalyzed H2A.Z deposition unravels the impact of nucleosome dynamics and the asymmetry of stepwise histone exchange

**DOI:** 10.1101/304998

**Authors:** Raushan K. Singh, Shinya Watanabe, Osman Bilsel, Craig L. Peterson

## Abstract

The SWR1C chromatin remodeling enzyme catalyzes an ATP-dependent replacement of nucleosomal H2A with the H2A.Z variant, regulating key DNA-mediated processes, such as transcription and DNA repair. Here we investigate the transient kinetic mechanism of the histone exchange reaction employing ensemble FRET, fluorescence correlation spectroscopy (FCS), and the steady state kinetics of ATP hydrolysis. Our studies indicate that SWR1C modulates nucleosome dynamics on both the millisecond and microsecond timescales, poising the nucleosome for the dimer exchange reaction. The transient kinetic analysis of the remodeling reaction performed under single turnover conditions unraveled a striking asymmetry in the ATP-dependent replacement of nucleosomal dimers, promoted by localized DNA translocation. Taken together, our transient kinetic studies identify new intermediates and provide crucial insights into the SWR1C-catalyzed dimer exchange reaction, as well as shedding light on how the mechanics of H2A.Z deposition might contribute to transcriptional regulation in vivo.

## INTRODUCTION

Eukaryotic genomes are assembled into long, linear arrays of nucleosomes that each consist of an octamer of core histones around which ∼147 bp of DNA is wrapped nearly two times. The histone octamer is composed of a central hetero-tetramer of histones H3 and H4, flanked by two heterodimers of histones H2A and H2B. Within the nucleosome, the H3/H4 tetramer wraps the central ∼90bp of nucleosomal DNA, while each H2A/H2B dimer organizes and stabilizes the final few turns (Luger *et al.*, 1997). In vivo, nucleosomal arrays are highly heterogeneous – nucleosomes are precisely positioned around regulatory regions, such as gene promoters or replication origins, different genomic regions harbor histones with a variety of posttranslational modifications, and the canonical core histones can be replaced with a number of conserved histone variants (Yuan et al., 2005; Raisner *et al.*, 2005; Jiang and Pugh, 2009; Vankatesh and Workman, 2015). These complex chromatin structures are often highly dynamic and can provide epigenetic information that regulates gene expression, replication timing, and other key nuclear processes (Swygert and Peterson, 2014; Henikoff, 2016)

Transcription in a eukaryotic cell is regulated by the structure and dynamics of nucleosomes located immediately upstream and downstream of the transcription start site (TSS) (Cairns, 2009; Dion et al., 2007). These promoter-proximal nucleosomes flank a nucleosome depleted region (NDR) of ∼200 bp, and are highly enriched for the conserved histone variant, H2A.Z (Hartley and Madhani, 2006; Barski et al., 2007). The H2A.Z variant is an evolutionarily conserved variant of H2A, whose incorporation into a nucleosome modulates its dynamics and promotes the intramolecular folding of nucleosomal arrays (Fan et al., 2002; Park et al., 2004). In budding yeast, H2A.Z is enriched in the promoter-regions of both active and inactive genes, and H2A.Z is known to play a key role in promoting the proper kinetics of transcriptional activation (Santisteban et al., 2000; Raisner *et al.*, 2005). In addition, yeast H2A.Z is enriched within nucleosomes that flank replication origins, as well as at the boundaries of heterochromatic regions where it mediates an anti-silencing effect by preventing the ectopic spread of heterochromatin (Albert et al., 2007; Meneghini and Madhani, 2003) Likewise, in higher metazoan, H2A.Z is enriched at pericentric heterochromatic regions during the early stages of embryonic development (Banaszynki et al., 2010). In addition to its critical role in transcription, H2A.Z has been intimately linked with DNA repair pathways and the regulation of cell cycle checkpoints, hallmarks of genome integrity (Adkins et al., 2013; Xu et al., 2012; Gevry et al, 2007). Not surprisingly, yeast cells lacking H2A.Z show temperature-sensitive growth defects and are sensitive to various genotoxic agents (Santisteban et al., 2000). Moreover, loss of H2A.Z in higher metazoans (such as frog and mouse) causes embryonic lethality (Faast et al., 2001).

Unlike canonical histones that are primarily assembled by a replication-dependent mechanism, H2A.Z is deposited at precise nucleosomal positions in an ATP-dependent reaction by enzymes related to the yeast SWR1C chromatin remodeling enzyme (Kobor et al., 2004; Mizuguchi et al., 2004). There are four subfamilies of chromatin remodeling enzymes, SWI/SNF, CHD, ISWI, and INO80, which are evolutionarily conserved from yeast to human (Clapier et al., 2017). Many chromatin remodelers are enormous, multi-subunit enzymes that each contain a related catalytic subunit that harbors a bi-lobular, RecA-like ATPase domain. SWR1C, and its mammalian paralogs, SRCAP and p400/Tip60, are members of the INO80C subfamily, and they use the energy of ATP hydrolysis to catalyze a histone exchange event where each of the two nucleosomal H2A/H2B dimers are sequentially replaced with H2A.Z/H2B variant dimers (Ruhl et al., 2006; Luk et al., 2010). Unlike all other chromatin remodelers that can use their ATP-dependent, DNA translocase activity to “slide” nucleosomes along DNA in cis, SWR1C can deposit H2A.Z without altering nucleosome positions (Clapier et al., 2017; Bowman 2010; Ranjan et al., 2015). In order to specifically direct the deposition of H2A.Z at promoter proximal nucleosomes, SWR1C is targeted to promoter regions by interactions with free DNA at the NDR, thereby targeting the adjacent +1 and −1 nucleosomes (nucleosomes are numbered relative to the transcription start site) (Ranjan et al., 2013). Likewise, the mammalian SRCAP and p400/Tip60 enzymes are believed to be targeted to promoter regions by gene-specific regulators (Pradhan et al., 2016; Yildirim et al., 2011).

The biological function of proteins requires local and global conformational fluctuations that take place in the micro– to millisecond time scale (Henzler-Wildman and Kern, 2007). Nucleosomes can undergo spontaneous conformational fluctuations in the millisecond time-scale that facilitate the transient accessibility of nucleosomal DNA to nuclear factors (Li and Widom 2004; Li et al., 2004). However, it remains unclear how remodeling enzymes, such as SWR1C, modulate the conformational dynamics of the nucleosome during an ATP-dependent nucleosome remodeling reaction. Notably, the SWR1C-catalyzed dimer-exhange reaction is complex, requiring the fine-coupling of the energy of ATP-hydrolysis to several microscopic events of the nucleosome remodeling reaction (Zhou et al., 2016). Therefore, this nucleosome remodeling cycle is expected to contain multiple intermediates which may remain invisible in discontinuous and/or steady-state biochemical assays. Transient kinetic experiments are well suited to unravel the indentity of reaction intermediates and microscopic rate constants associated with their production and decay, and hence provide in-depth mechanistic analysis of an enzyme-catalyzed reaction (Jencks 1987; Fersht 1999). Here we investigate the transient kinetic mechanism of the SWR1C-catalyzed dimer-exchange reaction employing ensemble FRET, FCS, and steady-state kinetics of ATP hydrolysis. We find that SWR1C utilizes an ATP-dependent modulation in nucleosome dynamics (in the microsecond time scale) as a strategy for discriminating the two structurally similar, H2A and H2A.Z nucleosomes. In addition, our FRET studies demonstrate that SWR1C uses a limited amount of DNA translocation to initiate the dimer exchange reaction, and that free H2A.Z/H2B dimers function as essential co-substrates that are required for resident H2A/H2B dimer eviction and subsequent replacement. Finally, our transient kinetic studies uncover asymmetry in the H2A.Z deposition reaction, where the linker distal dimer is replaced first, followed by the slower replacement of the linker proximal dimer. The asymmetry of the H2A.Z deposition reaction suggests a regulatory role for gene transcription, as well as providing insights into the molecular mechanism of ATP-dependent nucleosome remodeling catalyzed by other families of chromatin remodeling enzymes.

## RESULTS

### Dynamic nucleosome fluctuations specifies a substrate competent for SWR1C remodeling

To investigate the transient kinetic mechanism of SWR1C-catalyzed histone dimer exchange, a fluorescence-based strategy was employed. End-positioned, recombinant yeast mononucleosomes were reconstituted on a 202 bp fragment containing a “601” nucleosome positioning sequence. Nucleosomal substrates were designed with 55 bp of flanking free DNA so that it might reflect the asymmetry of a promoter proximal nucleosome located next to a nucleosome depleted region (NDR). The mononuclesome substrate contains a Cy3 fluorophore covalently attached to one end of the nucleosomal DNA and Cy5 attached to the H2A C-terminal domains (Figure 1A). These fluorophores are within an appropriate distance to function as a FRET pair, such that excitation of the Cy3 donor with a 530nm light source leads to efficient energy transfer to the Cy5 acceptor, as evidenced by the fluorescence emission peak at 670nm (Supplementary Figure 1).

**Figure 1.**
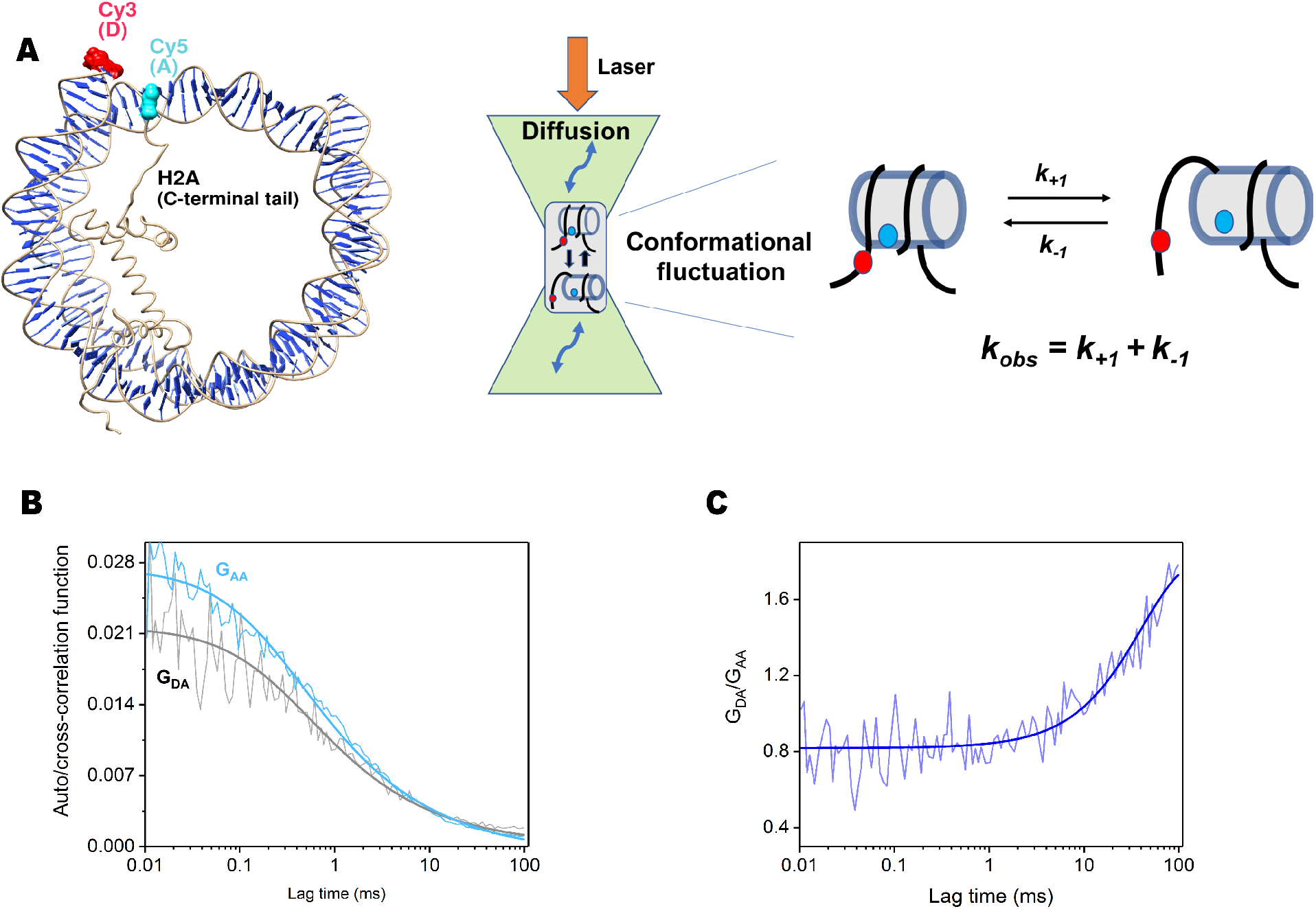
Measurement of conformational fluctuations of the nucleosome by FRET fluorescence correlation spectroscopy (FRET-FCS). **(A)** Experimental design. Left image shows the H2A-nucleosome substrate with Cy3 located on one DNA terminus and Cy5 attached to the H2A C-terminal domains (H2A-C120-Cy5). Middle image shows experimental set up for FCS-FRET. A femto-liter volume of nucleosome solution is excited by a laser at the donor excitation wavelength. Fluctuations in donor and acceptor fluorescence signals are due to two events: (1) diffusion in and out of the confocal volume and (2) nucleosome conformational fluctuations that are dictated by the intrinsic microscopic rate constants (*k_+1_* and *k_-1_*) causing a distance change between the donor-acceptor pair (right image). **(B)** The autocorrelation and cross-correlation functions of the acceptor and the donor-acceptor pair of the same nucleosome as a function of time are shown as cyan and black traces, respectively. **(C)** The ratio of the two correlation functions as function of time. The observed rate constant (*k_obs_*) of the conformational fluctuation is obtained from the exponential fit of the ratio curve of these two correlation functions.

Previous studies have demonstrated that nucleosomes undergo spontaneous unwrapping/rewrapping of nucleosomal DNA in the millisecond time scale (Li and Widom 2004; Li et al., 2004). To investigate the impact of SWR1C on this dynamic behavior, we investigated nucleosome dynamics utilizing FRET fluorescence correlation spectroscopy (FRET-FCS; Figure 1) (Torres and Levitus, 2007). In this assay, the conformational fluctuations of the nucleosome are determined from the ratio of the auto-correlation and cross-correlation functions of the change in fluorescence intensity of the acceptor (Cy5) and donor-acceptor (Cy3-Cy5) pair (Figure 1B, C). Utilizing FCS-FRET, the observed rate constant *(k_obs_*) for nucleosomal DNA unwrapping/rewrapping was determined to be ∼7 s^−1^ (half-life = ∼100 ms) (Figure 2A), slightly slower than values previously reported for a vertebrate nucleosome (∼21 s^−1^) (Li et al., 2004). Likewise, the dynamics of an H2A.Z nucleosome were similar, with a *k_ob_* of ∼2.1 s^−1^ (half-life =330 ms) (Figure 2D). Strikingly, binding of SWR1C to either an H2A or H2A.Z nucleosome increased the rate of DNA unwrapping/rewrapping by ∼2 orders of magnitude, compared to the unbound nucleosome (half-life = 1ms) (Figure 2B, E). Furthermore, addition of AMP-PNP further altered the dynamics of the SWR1C-H2A nucleosome complex, yielding a markedly biphasic pattern (Figure 2C). The two phases had nearly equal amplitudes, and the observed rate constants for the fast and slow phases were ∼40 s^−1^(half-life ∼1ms) and ∼5 ×10^4^ s^−1^(half-life ∼14 μs), respectively (Figure 2C). In contrast, addition of AMP-PNP had no impact on the dynamics of the SWR1C-H2AZ nucleosome complex (Figure 2F), suggesting that the enhanced microsecond dynamics are linked to substrate discrimination and may be key for subsequent catalytic steps.

**Figure 2.**
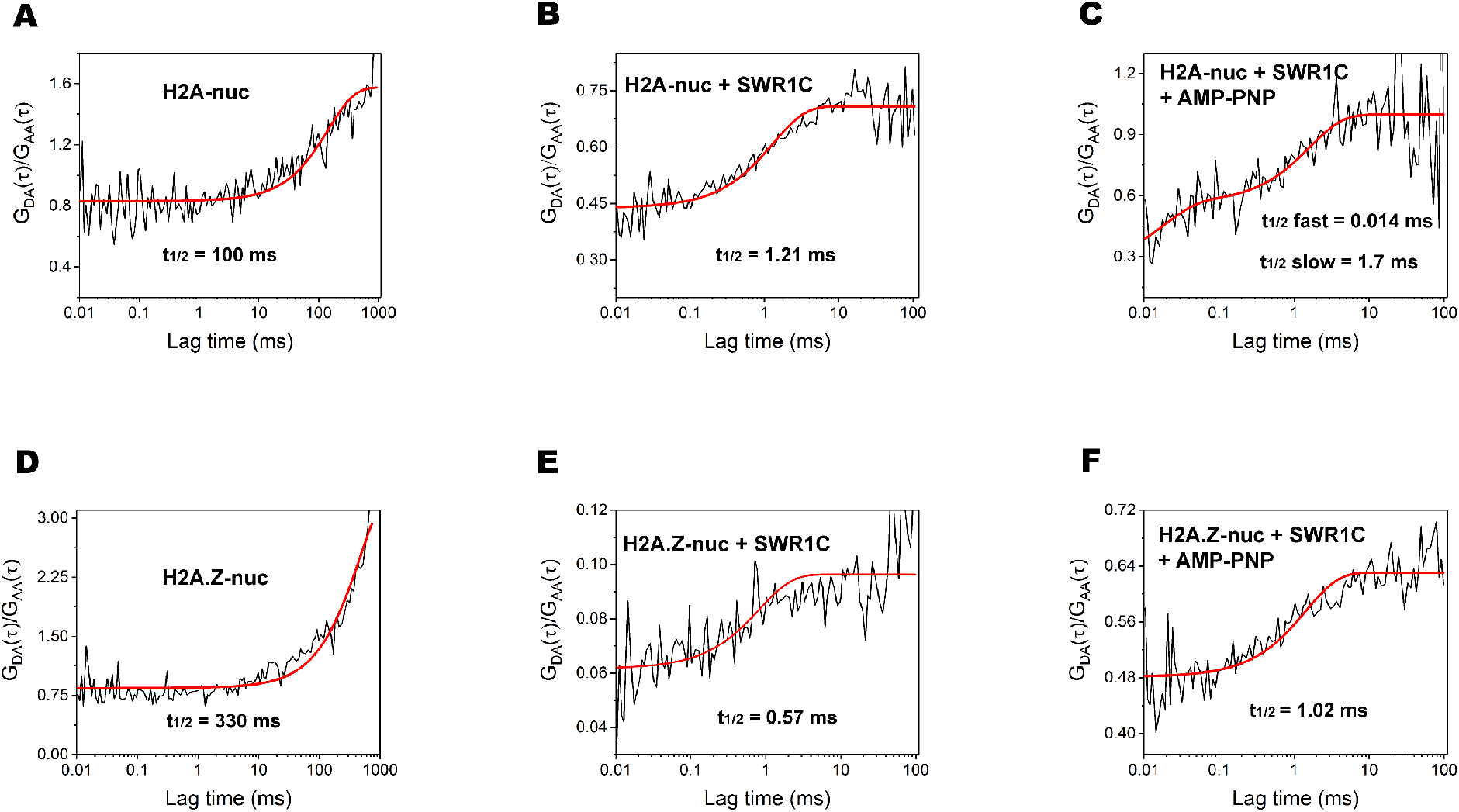
SWR1C modulates conformational fluctuations of the nucleosome. **(A)-(H)** The ratios of donor-acceptor cross-correlation to acceptor auto-correlation are plotted as a function of time under various experimental conditions marked with the associated panel. The experimental data were analyzed using either a single or double exponential rate equation yielding the values of the observed rate constants (t_1/2_ = 0.693/*k_obs_*) for the conformational fluctuation of the nucleosome. **(A)** Dynamics of an H2A-nucleosome; **(B)** Dynamics of the SWR1C-H2A nucleosome complex is 2 orders of magnitude faster than the free nucleosome; **(C)** Addition of AMP-PNP (a non-hydrolyzable analogue of ATP) to the SWR1C-nucleosome complex induces additional nucleosome dynamics on the microsecond timescale; **(D)** Dynamics of the H2A.Z nucleosome; **(E)** Dynamics of the SWR1C-H2A.Z nucleosome complex is 2 orders of magnitude faster than the free nucleosome; **(F)** Addition of AMP-PNP to the SWR1C-H2A.Z nucleosome does not alter nucleosome dynamics.

### SWR1C catalyzes ATP-dependent nucleosomal DNA translocation

A wealth of data support a unifying view that chromatin remodeling enzymes perform their various functions by initiating an ATP-dependent DNA translocation event from a fixed point on the nucleosome surface, about two DNA helical turns from the nucleosomal dyad (SHL +/-2.0) (Clapier et al., 2017). Indeed, SWR1C has been shown to make tight contact with nucleosomal DNA at SHL2.0, and single strand DNA gaps near SHL2.0 block H2A.Z deposition in vitro, suggesting an essential role for DNA translocation by SWR1C (Ranjan et al., 2015). However, unlike other remodeling enzymes, prior gel-based assays have not observed a translocation of DNA that leads to stable alterations in nucleosome positioning due to the SWR1C remodeling reaction (Luk et al., 2010; Ranjan et al., 2015). In order to directly probe for nucleosomal DNA translocation in real-time, we monitored the time-dependent loss of the nucleosomal FRET acceptor (Cy5) signal catalyzed by SWR1C under single-turnover conditions (excess enzyme to substrate). Importantly, addition of saturating amounts of SWR1C did not lead to a significant change in the Cy3 or Cy5 FRET signal of the nucleosome upon binding to SWR1C, suggesting that formation of the SWR1C-nucleosome complex does not unravel DNA from the nucleosomal edge (data not shown). Likewise, addition of a non-hydrolyzable ATP analog, AMP-PNP, did not lead to significant changes in the FRET signal (Figure 3A, red trace). In contrast, addition of ATP led to a time-dependent loss of the FRET signal, and analysis of the experimental data with a single exponential rate equation yielded the observed rate constant for nucleosomal DNA movement of 0.03 min^−1^ (Figure 3A, black trace). Notably, this slow loss of FRET signal was blocked by introduction of a pair of 2nt DNA gaps positioned at SHL-2.0 and SHL+2.0, consistent with ATP-dependent DNA translocation initiated near SHL2 (Figure 3B).

**Figure 3.**
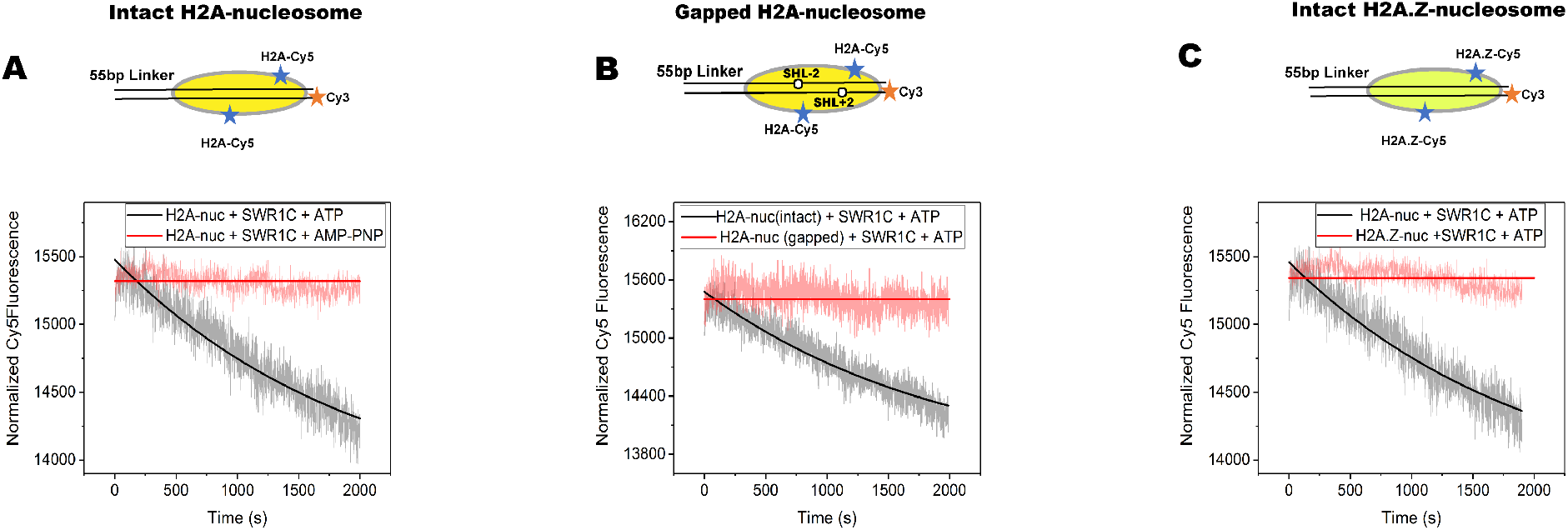
SWR1C catalyzes an ATP-dependent translocation of nucleosomal DNA. **(A)** The nucleosomal Cy5 FRET signal was monitored as a function of time in reactions that contained saturating amounts of SWR1C and ATP (black trace) or AMP-PNP (red trace). The kinetic trace for the reaction containing ATP was analyzed using a single exponential rate equation, yielding the value of *k_obs_* as 0.03±0.01 min^−1^. **(B)** Cy5 FRET signals for reactions contained SWR1C, ATP, and an intact FRET nucleosome (black trace) or a FRET nucleosome harboring 2 nt gaps at SHL+2.0 and SHL-2.0 (red trace). **(C)** Cy5 FRET signals for reactions containing SWR1C, ATP, and either an H2A (black trace) or an H2A.Z (red trace) nucleosome. All reactions shown were performed in parallel, and thus the reactions shown for the H2A nucleosome are from the same experiment. At least 3-4 kinetic traces were collected for each experimental condition and they were averaged. The resultant kinetic traces were analyzed using an exponential rate equation, and the error in the measurement represents the standard error of the parameter derived from non-linear regression analysis using Origin software package (OriginLab Corporation).

Furthermore, SWR1C was unable to catalyze an ATP-dependent loss of FRET on an H2A.Z nucleosome, supporting substrate specificity for DNA translocation (Figure 3C). This latter result was of particular significance given that SWR1C makes similar interactions and modulates the dynamics of an H2A.Z nucleosome in the absence of ATP, as assayed by FRET-FCS (Figure 2). We note, however, that DNA translocation activity correlates with the ability of SWR1C to induce an ATP-dependent, enhanced phase of nucleosomal dynamics on the microsecond timescale, suggesting that these rapid dynamics are linked to successful initiation of DNA translocation.

In parallel, we monitored DNA translocation by the related INO80C enzyme which repositions these nucleosomal substrates to a central location in an ATP-dependent reaction. In the presence of saturating levels of INO80C, a dramatic decrease in the Cy5 FRET signal was observed on both an H2A and an H2A.Z nucleosome, consistent with robust nucleosomal DNA translocation of >20 bp on both substrates (Supplementary Figure 2B, C). Furthermore, INO80C preferred the H2A.Z nucleosomal substrate, catalyzing a decrease in FRET at a ∼1.5-fold faster rate (∼0.6min^−1^ compared to 1.05min^−1^). Comparison of the amplitudes for the loss of Cy5 signal between the SWR1C and INO80C reactions indicates that DNA translocation in the SWR1C-catalyzed reaction is remarkably less than that of INO80C (Supplementary Figure 2A), suggesting that SWR1C may only translocate a few base pairs.

For remodelers such as INO80C that re-position nucleosomes from an end towards a center position, DNA is translocated from the linker proximal side of the nucleosome towards the linker distal side (Clapier et al., 2017), leading to extrusion of DNA and loss of the FRET signal when the FRET pair is located on a linker distal location. To test if SWR1C extrudes DNA from the distal nucleosomal edge or pulls DNA into the nucleosome, the Cy3 probe was incorporated at a location 15 bp within the nucleosome (Supplementary Figure S3A). On this substrate, addition of IN080C and ATP led to an initial increase in FRET signal, followed by a loss of FRET (Supplementary Figure S3C). This pattern is consistent with the initial movement of DNA towards the nucleosomal edge, followed by extrusion from the nucleosome. In contrast, addition of SWR1C and ATP did not lead to an increase in FRET signal, but rather yielded only a minor loss of FRET (Supplementary Figure S3B). These data suggest that SWR1C pulls a small amount of DNA into the linker distal side of the nucleosome.

### The SWR1C-catalyzed replacement of H2A-H2B dimers is markedly asymmetric

One consequence of localized DNA translocation might be the eviction of H2A/H2B dimers prior to replacement with H2A.Z/H2B. To monitor eviction of H2A/H2B dimers, an H2A-nucleosome was reconstituted that contained unlabeled nucleosomal DNA and the Cy3-Cy5 FRET pair located on the histone H3 N-terminus and the H2A C-terminus (Supplementary Figure 4A, middle schematic). Interestingly, addition of saturating concentrations of SWR1C and ATP did not decrease the FRET signal for this substrate, indicating that DNA translocation is not sufficient for dimer eviction (Supplementary Figure 4B). Previous studies have demonstrated that H2A.Z/H2B dimers function as co-substrates in the SWR1C exchange reaction, stimulating ATPase activity and interacting with both the Swr1 ATPase and the Swc2 subunit (Luk et al., 2010; Hong et al., 2014, Wu et al., 2005). Assayed under single turnover conditions, addition of free H2A.Z/H2B dimers to the SWR1C remodeling reaction led to a robust, extensive loss of Cy5 signal from the nucleosomal FRET substrate in an ATP-dependent reaction, indicating that H2A.Z/H2B dimers are essential co-factors for H2A/H2B eviction (Supplementary Figure 4C).

Eviction of H2A/H2B dimers was also monitored with the nucleosomal FRET substrate containing a Cy3-labelled DNA terminus that was used for the DNA translocation assays (Figure 4A). Whereas addition of SWR1C and ATP to this nucleosome catalyzed only a small decrease in the Cy5 FRET signal, further addition of free H2A.Z/H2B dimers led to a dramatic decrease in the FRET signal, consistent with eviction of the Cy5-labelled H2A/H2B dimers (Figure 4B). A comparison of the amplitude of the Cy5 FRET signal in reactions with and without H2A.Z-H2B dimers shows that addition of H2A.Z/H2B dimers causes an additional 10-fold decrease in the Cy5 FRET signal (Supplementary Figure 5). Notably, addition of free H2A/H2B dimers did not alter the Cy5 FRET signal, nor did H2A.Z/H2B dimers promote dimer loss from an H2A.Z nucleosome, results fully consistent with proper substrate specificity (Supplementary Figure 6). In addition, the H2A.Z/H2B-dependent loss of FRET was not observed on a substrate that contains a pair of 2nt DNA gaps at SHL+/-2.0, indicating that DNA translocation initiated near SHL2 is essential for dimer eviction (Figure 4E).

**Figure 4.**
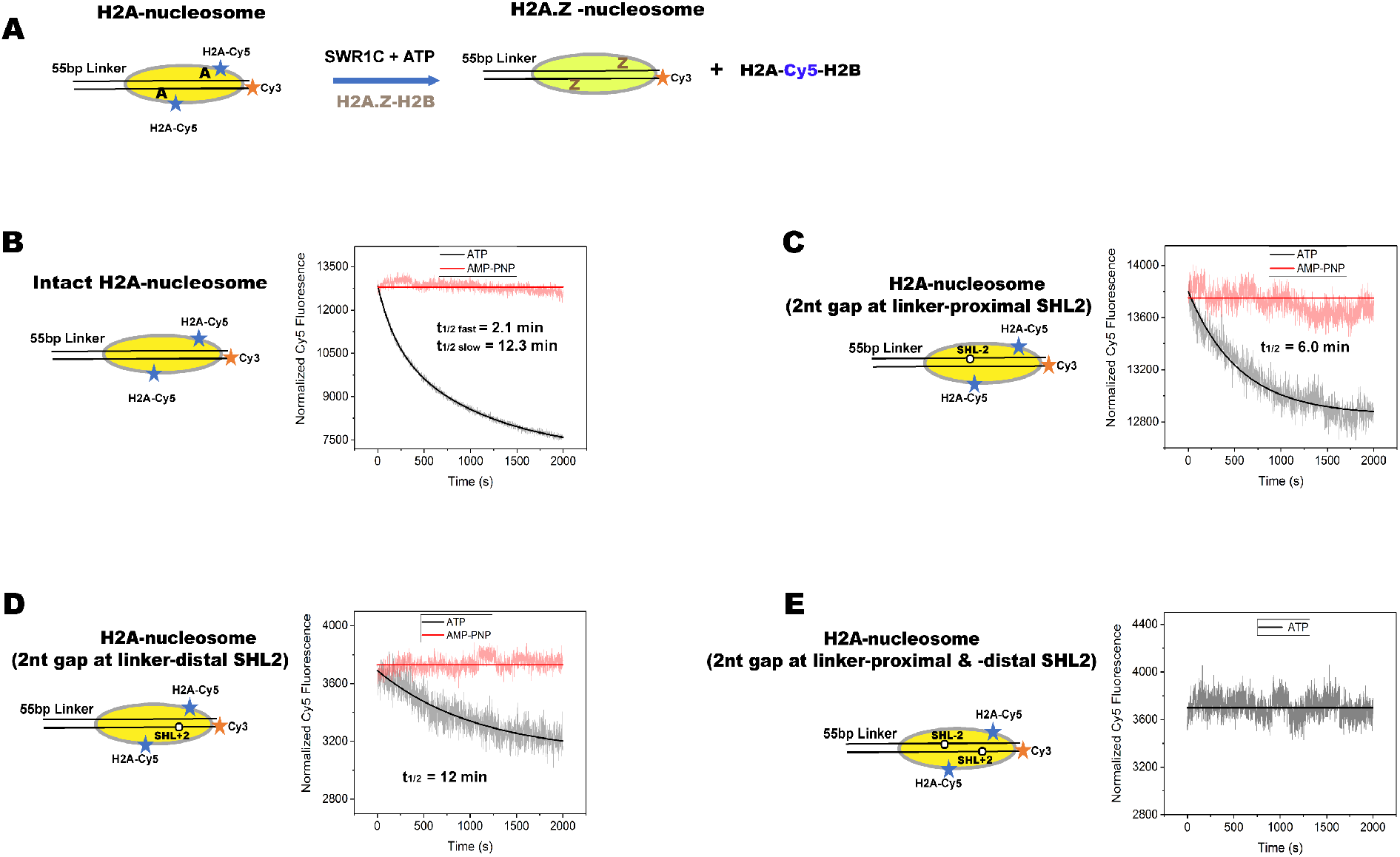
Transient kinetics of ATP-dependent eviction of two H2A-H2B dimers from an H2A-nucleosome is asymmetric. **(A-D)** Cy5 FRET signals were monitored over time as in Figure 3, but reactions also contained free H2A.Z/H2B dimers. **(A)** Schematic of reaction pathway. **(B)** Representative kinetic trace for SWR1C-catalyzed eviction of H2A-H2B dimers from an H2A nucleosome. The experimental data were analyzed using double exponential rate equation yield the observed rate constants (*k_obs_*) for the fast and slow phases as 0.33±0.02 min^−1^ (half-life = 2.1 min) and 0.06±0.01 min^−1^(half-life =12.3 min), respectively. **(B)** The kinetic trace for SWR1C-catalyzed eviction of H2A-H2B dimer from H2A-nucleosome harboring a 2nt gap at the linker-proximal SHL-2.0. The kinetic trace is monophasic, and hence it was analyzed using single exponential rate equation yielding the observed rate (*k_obs_*) as 0.12 ±0.03min^−1^(half-life = 6 min). **(C)** The kinetic trace for SWR1C-catalyzed eviction of H2A-H2B dimer from an H2A-nucleosome harboring a 2nt gap at the linker-distal SHL+2.0. The monophasic trace was analyzed using single exponential rate equation yielding the observed rate constant (*k_obs_*) as 0.06±0.01 min-^1^(half-life = 12min). **(D)** The kinetic trace for SWR1C-catalyzed eviction of H2A-H2B dimers from H2A-nucleosome containing 2nt gap at both SHL+2.0 and SHL-2.0. At least 3-4 kinetic traces were collected for each experimental condition and they were averaged. The resultant kinetic traces were analyzed using exponential rate equation, and the error in the measurement represents the standard error of the parameter derived from non-linear regression analysis using Origin software package (OriginLab Corporation).

Surprisingly, the kinetic trace of the dimer eviction reaction revealed a markedly biphasic reaction (Figure 4B). The experimental data were analyzed with a double exponential rate equation yielding values for the fast and slow observed rate constants (*k_obs_*) of 0.33 min^−1^ (halflife = 2.1 min) and 0.06 min^−1^ (half-life =12.3 min), respectively. In addition, the fast phase of the reaction was also associated with a large change in the FRET amplitude. One possibility is that the two, distinct kinetic phases reflect the sequential SWR1C-catalyzed eviction and replacement of each of the two H2A/H2B dimer under these single turnover conditions. To investigate this possibility further, we measured the kinetics for ATP-dependent deposition of H2A.Z-H2B. For monitoring H2A.Z deposition, the nucleosomal substrate contained a Cy3 fluorophore on the nucleosomal DNA edge, and the free H2A.Z/H2B dimer contained the Cy5 label on the H2A.Z C-terminus (Figure 5A). SWR1C reactions were initiated under single turnover conditions, and the kinetic trace shows an ATP-dependent increase in the FRET signal, consistent with H2A.Z deposition (Figure 5B). Importantly, the kinetic profile for H2A.Z deposition was also clearly biphasic, yielding observed rate constants (*k_obs_*) for the fast and slow phases of 0.32 min^−1^ (half life = 2.2) and 0.04 min^−1^ (half life = 16.6), respectively. Notably, the values of these observed rate constants for H2A.Z-H2B deposition are quantitatively similar to those of the fast and slow observed rate constants measured for the eviction of H2A-H2B (Figure 4). Taken together, the remarkable similarity in the biphasic kinetic profiles suggest that SWR1C catalyzes sequential exchange of two H2A-H2B dimers in a real-time assay performed under single turnover conditions.

**Figure 5.**
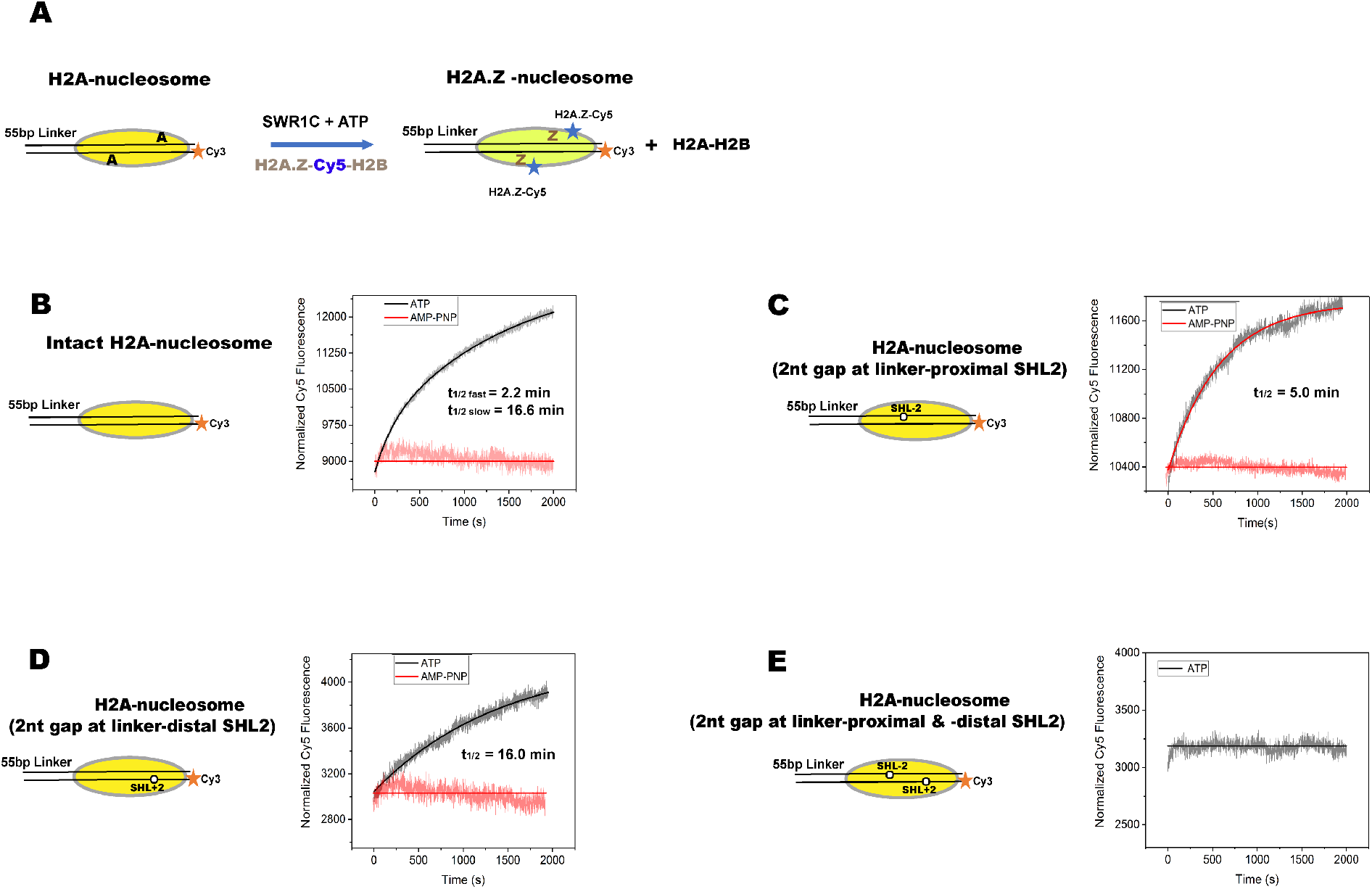
Transient kinetics of ATP-dependent deposition of two H2A.Z-H2B dimers is asymmetric. **(A)** Experimental strategy for monitoring the rate for deposition of H2A.Z-H2B. The nucleosomal substrate contains only the Cy3-labelled DNA end and Cy5 is located on the free H2A.Z/H2B dimer (H2A.Z-C126-Cy5). **(B)** Kinetic trace for the SWR1C-catalyzed deposition of H2A.Z-H2B dimer to the intact H2A-nucleosome. The biphasic trace was analyzed using double exponential rate equation yielding the observed rate constants (*k_obs_*) for the fast and slow phases as 0.31±0.01 min^−1^ (half-life = 2.2 min) and 0.04±0.01 min^−1^(half-life =16.6 min), respectively. **(C)** Reactions as in **(B)** but nucleosome harbors a 2nt gap at the linker proximal SHL-2.0. The monophasic trace was analyzed using single exponential rate equation yielding the observed rate (*k_obs_*) as 0.14±0.02 min^−1^(half-life = 5 min). **(D)** Same as in **(B)** but reactions contained a nucleosome with a 2nt gap at the linker-distal SHL+2.0. The monophasic trace was analyzed using a single exponential rate equation yielding the observed rate constant (*k_obs_*) as 0.04 ±0.01min-^1^(half-life = 16 min). **(E)** Reactions as in **(B)** but the nucleosome contained 2nt gaps at both SHL+2.0 and SHL-2.0. At least 3-4 kinetic traces were collected for each experimental condition and they were averaged. The resultant kinetic traces were analyzed using exponential rate equation, and the error in the measurement represents the standard error of the parameter derived from non-linear regression analysis using Origin software package (OriginLab Corporation).

The biphasic kinetics of dimer eviction and deposition may reflect asymmetry in the catalytic cycle, such that the first round of dimer exchange occurs preferentially on one face of the nucleosome with a rate that is ∼6-fold faster than exchange of the second dimer. To address this question, FRET mononucleosomes were reconstituted that contained single, 2nt gaps in nucleosomal DNA at either the linker-proximal (SHL-2.0) or linker-distal (SHL+2.0) regions (Figure 4C,D). These substrates were used in dimer eviction reactions peformed under single turnover condition with SWR1C, H2A.Z/H2B dimers, and ATP. Notably, the ATP-dependent kinetic profiles for these gapped substrates were monophasic—in sharp contrast to the mononucleosome with intact nucleosomal DNA. For instance, when the 2nt gap was located at linker distal SHL+2.0, only the slow phase (*k_obs_* = 0.06 min^−1^) of FRET loss was observed, and the change in FRET amplitude was small (Figure 4D). In contrast, when the gap was located at the linker proximal SHL-2.0, only the fast phase (*k_obs_* = 0.12 min^−1^) of FRET loss remained (Figure 4C). Furthermore, this fast phase was associated with a much larger drop in the FRET amplitude, compared to the slow phase, indicating that the fast phase reflects removal of the dimer closest to the distal, Cy3-labeled DNA. Together, these results indicate that SWR1C preferentially evicts and replaces the H2A-H2B dimer located at the linker-distal half of the nucleosome, followed by a slower reaction where the linker-proximal H2A/H2B dimer is replaced.

The impact of single 2nt gaps on the H2A.Z deposition reaction was also monitored (Figure 5). As expected from the dimer eviction data, the kinetic traces for these reactions with substrates containing single 2nt gaps were monophasic (Figure 5C,D), in contrast to reactions using an intact nucleosome (Figure 5B). Furthermore, the observed rate constants and the associated amplitudes for H2A.Z-H2B deposition with nucleosomal substrates with gaps at either SHL-2.0 or SHL+2.0 were quite similar to the observed rate constants for the fast and slow kinetic phases of dimer deposition on the intact nucleosome (Figure 5B-D). Taken together, the observed rate constants for the sequential exchange of both H2A-H2B dimers unravels marked spatio-temporal asymmetry of the SWR1C-catalyzed H2A.Z deposition cycle.

### SWR1C-nucleosome interactions couple ATPase activity to DNA translocation

Remodeling enzymes couple the energy of ATP hydrolysis to translocation of DNA at SHL2.0 of the nucleosome, and gaps at SHL2.0 block DNA translocation and dimer exchange (Figure 3 and Ranjan et al., 2015). Previous studies have shown that the basal ATPase activity of SWR1C is stimulated by both the nucleosomal substrate and the H2A.Z./H2B co-substrate (Figure 6A; Luk et al., 2010). To probe the impact of intact nucleosomal DNA on the chemo-mechanical coupling of SWR1C ATPase activity, steady-state ATPase assays were performed with a nucleosomal substrate that contains 2nt gaps at both SHL+2.0 and SHL-2.0 (Figure 6B,C). Strikingly, the gapped nucleosome was unable to stimulate the ATPase activity ofSWR1C (Figure 6B). Thus, the stimulation of SWR1C ATPase activity by nucleosomes reflects efficient coupling of ATP hydrolysis to productive interactions with DNA that lead to translocation. In sharp contrast, gaps in nucleosomal DNA did not diminish the impact of H2AZ/H2B, but led to a further ∼1.5x increase in the steady state rate (Figure 6B). Thus, on a gapped nucleosome, the H2AZ/H2B dimers stimulate the rate of hydrolysis, reflecting an uncoupling of ATP hydrolysis from DNA translocation. The impact of the gap appears similar to the ATPase cycle of AAA+ chaperones which undergo rapid hydrolysis of ATP upon encountering a very stable substrate that is resistant to ATP-dependent unfolding (Sauer and Baker, 2011).

**Figure 6.**
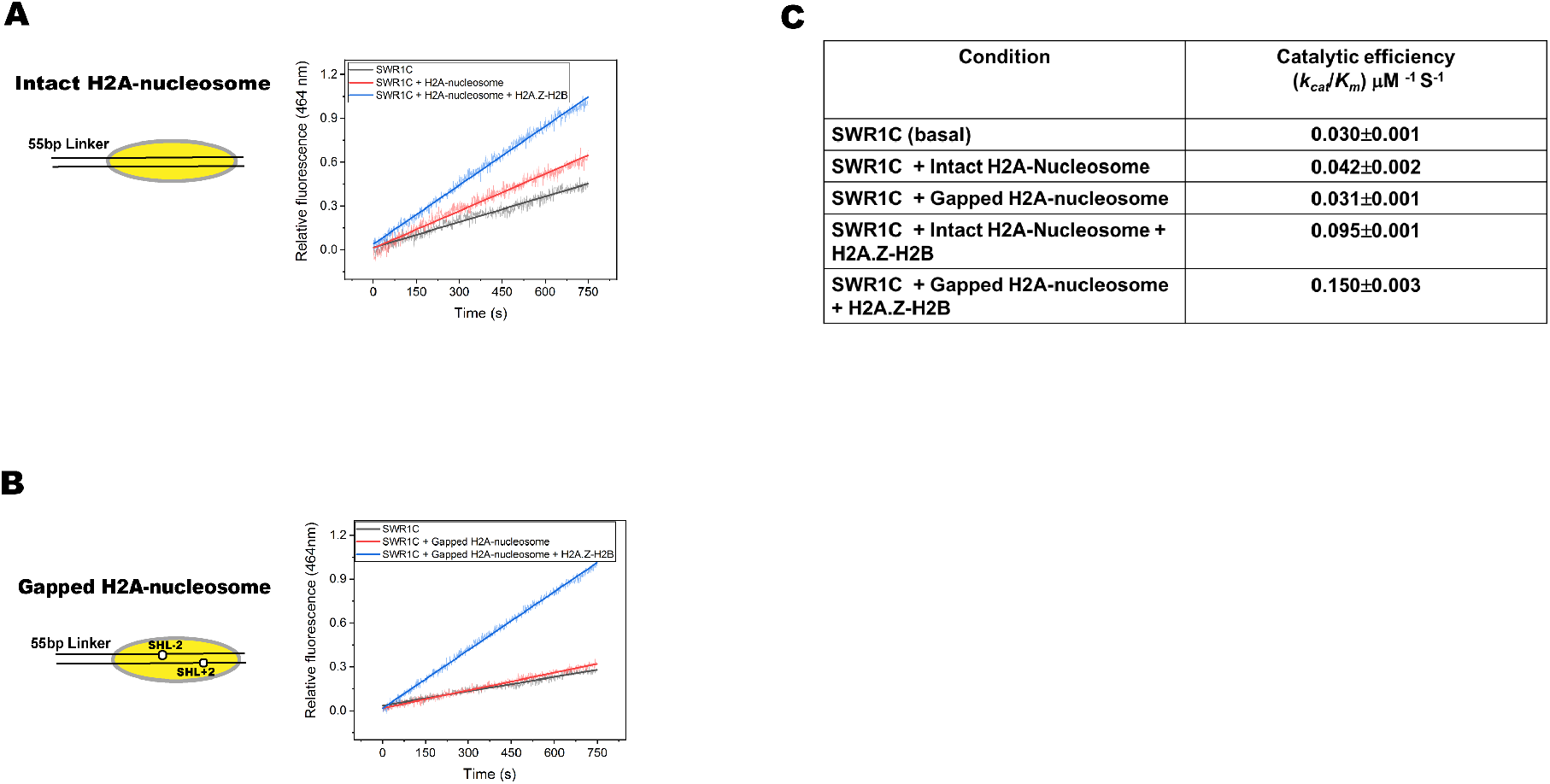
The presence of 2nt gaps at the SHL±2 region of nucleosomal DNA disrupt ATPase coupling. **(A)** Steady state kinetics of SWR1C ATPase activity. The black and red traces show ATPase activity of SWR1C in the absence (basal), or presence of an H2A nucleosome (intact nucleosome), respectively. The blue trace shows ATPase activity in the presence of an H2A nucleosome and free H2A.Z/H2B dimers. **(B)** Reactions as in **(A)** but the nucleosomal substrate contains 2nt gaps at both SHL-2.0 and SHL+2.0 (gapped nucleosome). At least three kinetic traces were collected under the above experimental conditions. The averaged kinetic traces were analyzed using the steady-state rate equation as described in the method section. **(C)** The values of *k_cat_/K_m_* for SWR1C ATPase activity which represent the substrate specificity of the enzyme for ATP in the presence of the its co-substrates (H2A-nucleosome and H2A.Z-H2B dimer) under the above experimental conditions are listed in this table.

## Discussion

SWR1C is unique among remodeling enyzmes as it cannot mobilize nucleosomes in *cis*, but rather it is dedicated to the ATP-dependent replacement of nucleosomal H2A with its variant, H2A.Z (Clapier et al., 2017). In contrast to previous studies of ATP-dependent, nucleosome sliding reactions, we found that the dimer exchange reaction is kinetically slow, likely because the reaction has to transit multiple activation or transition state barriers during the catalytic cycle (Hammes, 2002). Furthermore, the coordination of several different microscopic events associated with each round of dimer exchange – DNA translocation, H2A-H2B eviction, and H2A.Z-H2B deposition – is likely to yield a large number of kinetic intermediates. Using multiple biophysical approaches, our transient kinetic investigation supports a complex reaction pathway, involving at least five distinct intermediates (Figure 7): (Step 1) Binding of SWR1C to an end-positioned, asymmetric nucleosome yields a SWR1C-nucleosome complex that has a ∼100-fold enhanced rate of DNA wrapping/unwrapping; (Step 2) Binding of ATP leads to an additional enhancement of nucleosome dynamics on the microsecond timescale that are unique to an H2A nucleosomal substrate; (Step 3) ATP-hydrolysis promotes a small amount of DNA translocation from SHL+2.0 that is not sufficient for H2A/H2B eviction; (Step 4) free H2A.Z/H2B dimers act as a power stroke, promoting ATP hydrolysis, leading to the initial eviction of the linker-distal H2A/H2B dimer and replacement by H2A.Z/H2B in an apparently concerted reaction; (Step 5) The second, linker-proximal dimer is sequentially replaced during the single turnover reaction cycle with kinetics ∼6-fold slower than the first replacement event. Below, we discussed in greater detail the mechanistic implications for this reaction series.

**Figure 7.**
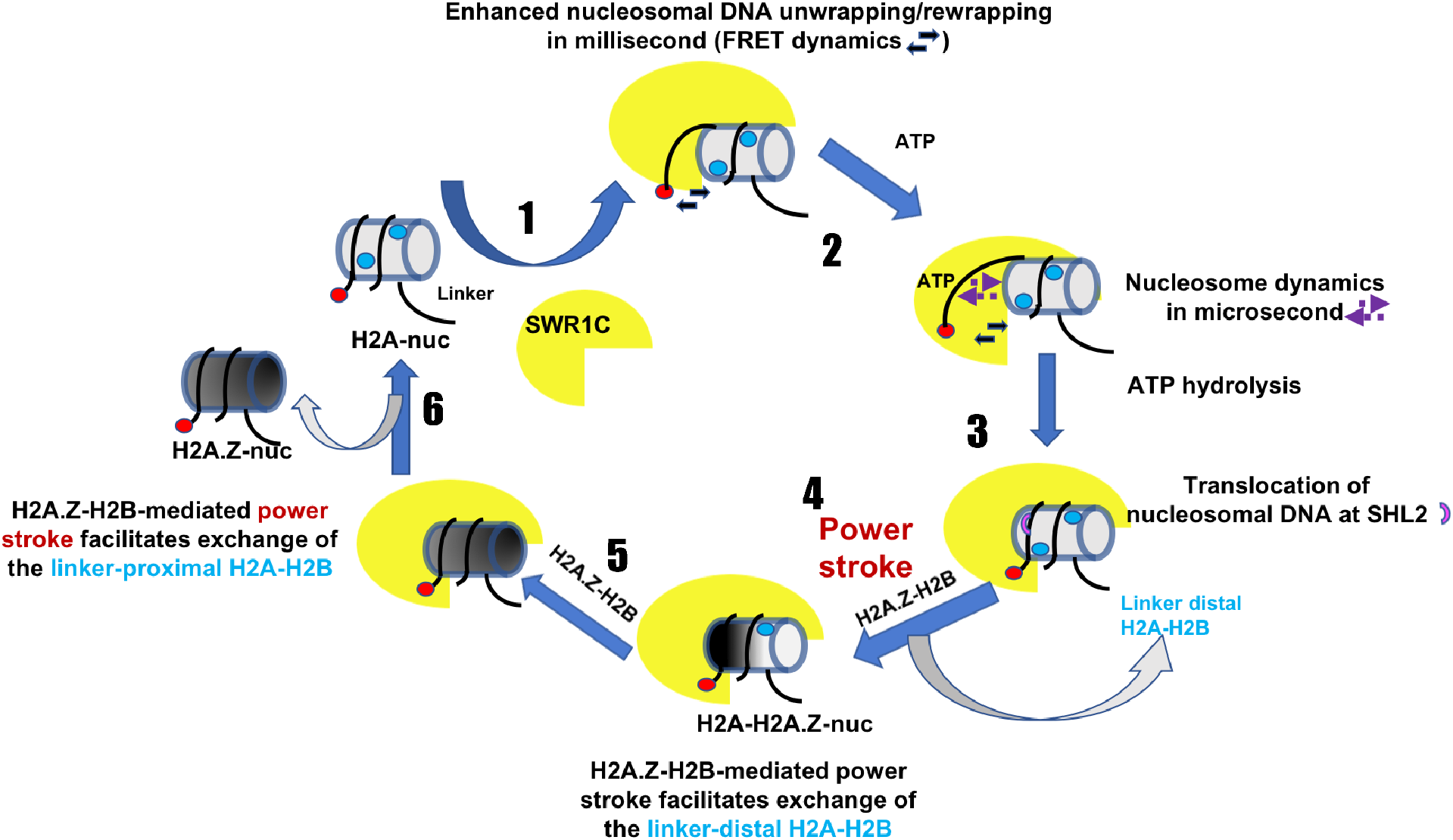
Kinetic model for the SWR1C-catalyzed histone dimer exchange reaction. **(1)** The engagement of SWR1C to the H2A-nucleosome enhances the unwrapping/rewrapping kinetics of the nucleosomal DNA in the millisecond time scale. **(2)** Binding of ATP to the SWR1C engaged nucleosome further impacts its dynamics in the microsecond time scale. **(3)** SWR1C catalyzes a slow translocation of nucleosomal DNA, coupled with ATP hydrolysis. **(4)** Binding to H2A.Z-H2B to SWR1C activates ATPase activity, and is proposed to generate a power stroke required for the replacement of linker-distal dimer. **(5)** SWR1C remains engaged with the H2A-H2A.Z heterotypic nucleosome and catalyzes the slower replacement of the linker-proximal H2A-H2B dimer, utilizing the H2A.Z-H2B-mediated second round of power stroke.

### Conformational fluctuations of the nucleosome during the dimer exchange reaction

Macromolecules undergo spontaneous conformational fluctuations, leading to ensembles of multiple, distinct conformations (Henzler-Wildman and Kern, 2007). Notably, biophysical studies have shown that such “wiggling and giggling” in proteins or enzymes are indispensable for their function, and that these dynamics often impact enzyme-substrate specificity and are kinetically coupled with their catalytic turnover rate (Feynman et al, 1963, Agarwal et al, 2002; Henzler-Wildman et al, 2007). The nucleosome is known to undergo spontaneous conformational fluctuations in the millisecond time scale, manifested in the unwrapping and rewrapping of nucleosomal DNA (Li and Widom 2004; Tims et al, 2011). Additional conformational fluctuations are also likely to involve the entire nucleosome (Henzler-Wildman and Kern, 2007), including the histone octamer, and such dynamics are expected to influence remodeling reactions.

We found that the binding of SWR1C to the nucleosome is characterized by a ∼100-fold increase in the rate of nucleosome conformational fluctuations in the millisecond timescale. Faster unwrapping/rewrapping kinetics of the nucleosomal DNA end is likely to facilitate the eviction of H2A-H2B dimers, as the dimers are tightly held within the nucleosome via a strong electrostatic interaction with the last 3 superhelical turns (SHL+/-3.5-6.5) of nucleosomal DNA (Luger et al., 1997). Additionally, these conformational fluctuations may also promote the generation of early intermediates of the dimer-exchange reaction by reducing the activation energy barrier for approaching the transition state (Daniel et al 2003; Nashine et al, 2010). This viewpoint is strengthened by our observation that ATP binding induces additional nucleosomal fluctuations in the microsecond time scale, changes that are not observed when SWR1C is bound to the remodeling product, the H2A.Z nucleosome. Such a stark difference in the conformational fluctuations between an H2A and H2A.Z nucleosome underscores the idea that kinetic coupling of nucleosomal conformational fluctuations may be critical for progression of the ATP-dependent dimer exchange reaction cycle (Eisenmesser et al, 2002). We also envision that ATP-dependent nucleosome dynamics may facilitate the ability of SWR1C to search for an appropriate conformation of nucleosome to be funneled into the catalytic cycle (Vendruscolo and Dobson 2006). Notably, the catalytic efficiency of an enzyme is often linked with the kinetics of a conformational search of both the enzyme and its cognate substrate (Benkovic and Hammes-Schiffer 2003). Thus, in this view, the ATP-bound, SWR1C-H2A.Z-nucleosome complex may be kinetically trapped at the beginning of the catalytic cycle.

Recently, two studies have reported cryo-EM reconstructions of the yeast and human IN080C remodeling enzyme bound to an end-positioned nucleosome (Eustermann et al., 2018; Ayala et al., 2018). IN080C is highly related to SWR1C, having a similar subunit module organization and sharing several subunits, such as the Rvb1/Rvb2 heterohexomeric ring assembly. Remarkable, INO80C and SWR1C appear to use different strategies to engage the nucleosome. First, INO80C interacts with nearly an entire gyre of nucleosomal DNA on the linker proximal side of the nucleosome, and the ATPase lobes of the Ino80 subunit make tight contact with the linker proximal SHL-6 region rather than at SHL-2, consistent with previous footprinting studies (Brahma et al., 2017). These interactions position IN080C to initiate DNA translocation from the linker proximal side, pulling the long linker DNA into the nucleosome and re-positioning the nucleosome towards the center of the DNA fragment. The authors also suggest that interactions on this nucleosomal face may also facilitate exchange of the linker proximal dimer. Interestingly, previous hydroxyl radical footprinting studies have shown that SWR1C also makes contact with this face of the nucleosome, interacting with the long linker DNA and weakly contacting SHL-5.5 (Ranjan et al., 2015). However, SWR1C does not appear to contact this entire gyre of DNA, but rather SWR1C makes the strongest contacts at the linker distal SHL+1.0 and +2.0 region. Thus, SWR1C appears to interact across both gyres of DNA, positioning itself to initiate DNA translocation from the linker distal side of the nucleosome, promoting the first round of dimer exchange. Interactions with linker DNA may help to recruit or orient SWR1C, or such contacts may prevent propagation of the DNA translocation event such that nucleosome positions are unchanged (Clapier et al., 2017).

Remarkably, binding of IN080C to the nucleosome releases ∼15bp of DNA from the histone octamer surface where the ATPase lobes interact at SHL-6 (Eustermann et al., 2018; Ayala et al., 2018). In contrast, nucleosome binding by SWR1C does not appear to stably disrupt histone-DNA contacts at the distal nucleosome edge, as we detect no change in steady-state FRET when probing either the DNA terminus or 15bp inside the nucleosome. However, the SWR1C-nucleosome complex does show greatly enhanced dynamics of nucleosomal DNA. Such rapid unwrapping/rewrapping events may be analogous to a “snapshot” of an unraveled state that may be trapped in the cryogenic INO80C-nucleosome complex.

### Nucleosomal DNA translocation is critical for the histone replacement reaction

SWR1C shares two common features with other remodeling enzymes: it makes tight contacts with nucleosomal DNA at nucleosomal location SHL2, and its remodeling activity is blocked by single-strand gaps on the 3’ to 5’ DNA strand, implicating DNA translocation as an essential step (Ranjan et al., 2015; Bartholomew 2014). However, in the case of SWR1C, only gaps in very close proximity to its binding site at SHL2 block dimer exchange (+/-17bp to +/-23bp from the nucleosomal dyad) (Ranjan et al., 2015), consistent with a requirement for <6bp of DNA translocation. Indeed, our ensemble FRET assays suggest that only a few bps of DNA are translocated by SWR1C following addition of ATP. Notably, the observed rate constant for this DNA translocation step is several fold slower than that of the subsequent H2A/H2B dimer eviction step which is dependent on addition of the free, H2A.Z/H2B cosubstrate. We propose that H2A.Z/H2B enhances the rate, but not the extent of nucleosomal DNA translocation. Such rapid, but limited amount of DNA translocation, may not only pull DNA into the nucleosome, but also lead to allosteric changes in the histone octamer that destabilize the H2A/H2B and H3/H4 interface (Sinha et al., 2017).

### Chemo-mechanical coupling in the ATP-dependent nucleosome remodeling reaction

The H2A.Z/H2B co-substrate binds to the Swr1 ATPase in a region N-terminal to the ATPase lobes. This region is adjacent to the HSA domain that binds to an Actin/Actin-related protein (ARP) subunit module (Wu et al, 2009; Nguyen et al, 2013). Recent studies with the RSC remodeling enzyme have shown that a similar ARP subunit module also recognizes the HSA domain of the Sth1 ATPase, and that these subunits directly interact with the ATPase lobes to stimulate ATP hydrolysis and DNA translocation. Importantly, this activation process is key for RSC to promote nucleosome disassembly (Clapier et al., 2016). We propose that the binding of the H2A.Z/H2B cosubstrate to Swr1 may also promote interactions between the adjacently bound, Actin/Arp module and the ATPase lobes, and that this mechanism provides an explanation for the activation of ATPase activity. Like the case with RSC, such stimulation of the ATPase and DNA translocation activities of SWR1C would promote nucleosome disassembly (e.g. dimer eviction).

Nucleosome binding enhances the affinity for ATP and stimulates the rate of ATP hydrolysis (Luk et al., 2010). How could binding of SWR1C to the nucleosome enhance the rate of ATP hydrolysis? In the SWI2/SNF2 ATPase, DNA-stimulated ATPase activity has been attributed to a DNA-mediated rearrangement of the ATPase lobes that orients catalytic residues for ATP hydrolysis (Durr et al., 2005). Likewise, a recent cryo-EM structure of the Chd1-nucleosome complex shows that the two ATPase lobes of the remodeler undergo a well pronounced structural change in the presence of a ground state analogue of ATP (ADP-BeF3) (Farnung et al., 2017), inducing close interactions with the nucleosome at the SHL2 region. We suggest that the Swr1 ATPase may undergo a similar rearrangement upon binding to nucleosomal DNA at SHL±2 that poises it for rapid hydrolysis. Consistent with this view, our studies demonstrate that the stimulation of ATP hydrolysis is eliminated by a 2nt gap at SHL2, indicating that tracking of nucleosomal DNA is fine-tuned with the kinetic events of the ATPase cycle. Thus, intact nucleosomal DNA is likely to provide a macro-molecular context essential for optimum closure of the ATPase lobes upon the ATP binding (Durr et al., 2005; Farnung et al., 2017).

### SWR1C catalyzes an asymmetric dimer exchange reaction

Previous gel-based assays for H2A.Z deposition demonstrated that the dimer exchange reaction is a sequential (Luk et al., 2010), step-wise process when assayed under steady state assay conditions (excess substrate to enzyme). We were surprised, however, to find that SWR1C catalyzes two, sequential rounds of dimer exchange even under single turnover reaction conditions (excess enzyme to substrate). Our single turnover reactions were clearly biphasic, with the first phase occurring at a rate about ∼6-fold faster than the second phase. Furthermore, 2nt DNA gaps at either SHL+2.0 or SHL-2.0 produced monophasic kinetic profiles that maintained either the fast or slow rates observed with intact nucleosomes (Figure 4). These data suggest the intriguing possibility that SWR1C catalyzes two sequential rounds of dimer exchange without a requisite dissociation from the nucleosome substrate. Furthermore, the slower rate of the second phase suggests that the second round of dimer exchange has a different rate-limiting step or has an altered reaction pathway.

How might SWR1C accomplish this feat? We envision that following exchange of the first H2A/H2B dimer, SWR1C must re-orient its ATPase lobes to the adjacent DNA gyre so that it can initiate a DNA translocation event that “pulls” the long linker DNA end into the nucleosome, promoting eviction of the linker proximal dimer. Importantly, re-orientation of the lobes would not require dissociation of the entire enzyme from the nucleosome. In this model, the ATPase lobes may interact with the neighboring SHL-2.0, or it may involve binding to SHL-5.5 which is directly adjacent to its initial binding site at SHL+2.0. Intriguingly, a recent study has suggested that the Chd1 remodeling enzyme may re-orient its ATPase lobes back and forth between SHL2.0 and SHL5.5 during ATP-dependent nucleosome mobilization (Qiu et al., 2017). Indeed, a recent cryo-EM structure of a Chd1-nucleosome complex shows binding of the ATPase lobes to SHL5.5 (Farnung et al., 2017). In addition, the related INO80C ATPase is known to catalyze nucleosome sliding by using interactions between its ATPase lobes and SHL5.5 on the nucleosome (Brahma et al., 2017; Eustermann et al., 2018; Ayala et al., 2018). Flexibility of the remodeler ATPase lobes for multiple, alternative interactions with nucleosomal DNA may be a hallmark of these enzymes.

From yeast to mammals, H2A.Z deposition appears to be targeted to the nucleosome adjacent to the start site for transcription by RNA polymerase II (Albert et al., 2007; Barski et al., 2007). Often termed the +1 nucleosome, it is inherently asymmetric, with one side flanked by a nucleosome depleted region (NDR) of 140-250 bp, and the other side by the +2 nucleosome which can be separated from the +1 by less than 20 bp of linker DNA (Jiang and Pugh 2009). In yeast, targeting of SWR1C to the +1 nucleosome relies on protein-DNA interactions between SWR1C and the NDR region (Ranjan et al., 2013), whereas the related vertebrate enzymes, SRCAP and p400/Tip60, are believed to be recruited to promoter proximal regions by gene-specific regulators (Pradhan et al., 2016). Our in vitro nucleosome substrate mimics the asymmetry of the +1 nucleosome, as it is flanked by a 55 bp linker DNA. Interactions between SWR1C with the long linker DNA appears to orient the ATPase lobes of the Swr1 catalytic subunit to interact with linker distal SHL+2.0, leading to preferential replacement of the linker distal H2A/H2B dimer in the initial, fast phase of the biphasic exchange reaction (Figure 4B). Recent high resolution ChIP-exo analyses of nucleosome asymmetry in yeast are fully consistent with asymmetric dimer exchange (Rhee et al., 2014). At the +1 nucleosome, the promoter distal half of the nucleosome is highly enriched for H2A.Z, whereas the promoter proximal side is enriched for H2A. Interestingly, the promoter proximal side is also enriched for ubiquitinylated H2B (H2B-ub), a mark associated with active transcription (Rhee et al., 2014; Zhang 2003). One interesting possibility is that H2B-ub might enhance the intrinsic kinetic delay of the second round of dimer exchange, ensuring that the +1 nucleosome remains asymmetric with respect to H2A.Z deposition.

What might be the functional significance of dimer exchange asymmetry? We consider two possibilities that would be consistent with the known role of H2A.Z in promoting rapid induction of transcription from a poised promoter (Guillemette et al., 2005). First, there may be unique biochemical properties for a heterotypic H2A.Z/H2A nucleosome, especially when the H2A/H2B dimer contains a mono-ubiquitin mark. H2B-ub can disrupt nucleosome-nucleosome interactions in vitro (Fierz et al., 2011), and together with H2A.Z, this combination may favor subsequent nucleosome disruption during transcription initiation. Alternatively, the kinetic lag between the first and second rounds of dimer exchange may lead to an accumulation of a remodeling intermediate where SWR1C enhances the wrapping/unwrapping dynamics of nucleosomal DNA on the NDR-proximal side. In yeast, the NDR proximal side of the nucleosome often contains the site of transcription initiation (Jiang and Pugh 2009), and thus a mechanism that specifically enhances accessibility to this face of the nucleosome would be particularly advantageous.

## Acknowledgements

We thank Nate Giaocchini (UMMS) for optimizing the purification of SWR1C and other members of the Peterson lab for helpful discussions. We also thank David Lambright (UMMS) for access to his ISS PC1 spectrofluorometer. This work was supported by a grant from the NIH (GM049650) to C.L.P. and a postdoctoral fellowship from the American Heart Association to R.K.S.

## Author Contributions

All experiments were performed by R.K.S.; S.W. provided purified IN080C; O.B. provided assistance with the FCS studies; C.L.P. and R.K.S. analyzed the data and prepared the manuscript.

## Experimental Procedures

### Reconstitution of fluorescently labeled mononucleosomes

Recombinant yeast histones (H2A, H2B, H3 and H4) were expressed in *E. coli* and purified from inclusion bodies as described previously (Luger et al., 1999). The unique cysteine substitutions were introduced at H2A-120 and H3-33 using quick-change mutagenesis. Histones were labeled with Cy5 and Cy3 using maleimide chemistry as described previously. The 202 bp Cy3-labeled DNA fragment that contains an end positioned 601 nucleosome positioning sequence was prepared by PCR amplification utilizing the end-labeled or internal-labeled PCR-primers purchased, respectively, from IDT and IBA. Fluorescent mononucleosomes were reconstituted by salt dialysis (Luger et al., 1999). For each set of reconstitutions, at least three different ratios of histone octamer to DNA template were assembled, and the reconstitution that yielded 1-5% free DNA was chosen for subsequent reactions. The gapped mononucleosomes were reconstituted using the 202 bp DNA fragment containing end positioned 601-positioning sequence harboring 2nt gap at the SHL±2 region. The gapped DNA fragment was generated by PCR amplification utilizing the primers that contain deoxyuridine residues at the specific gap sites. In order to create a gap in the above PCR product, it was treated with USER enzyme (NEB)— a mixture of DNA glycosylase and endonuclease III. The completion of the deoxyuridine removal from the PCR product (by USER enzyme) was confirmed upon its treatment with S1 nuclease (Thermo Fisher Scientific).

### Purification of yeast SWR1C

SWR1C was purified from whole cell extracts of a yeast strain harboring a FLAG-tagged allele of the Swr1 ATPase (Swr1-3xFLAG) as detailed elsewhere (Mizuguchi et al., 2012). Following FLAG peptide elution, SWR1C was further purified on a 5ml, 5-30% glycerol gradient in buffer D (25mM HEPES, pH = 7.6, 1mM EDTA, 2mM MgCl_2_, 100mM KCl). Gradients were sedimented for 14 hours at 35000 RPM. Peak fractions of SWR1C were pooled, concentrated using centricon (10kDa cutoff), and dialyzed overnight against storage buffer (25mM HEPES, pH = 7.6, 1mM EDTA, 2mM MgCl_2_, 100mM KCl, and 10% glycerol). SWR1C was flash frozen and stored at −80°C until use. SWR1C concentration was determined by SDS PAGE analysis along side a BSA standard titration, followed by Sypro staining and quantification on a Typhoon imager.

### Nucleosome dynamics measurements using fluorescence correlation spectroscopy

FCS measurements were carried out using an in-house automated FCS set up. Excitation was provided by a 488 nm single-mode fiber coupled picosecond diode laser (BDL-488-SMN Becker & Hickl GmBH) that was expanded to overfill the microscope objective. The excitation was focused on the sample and collected by an Olympus UPlanSApo 60x 1.2 N.A. water immersion objective. The collected light was focused on a 50 micron pinhole and then further collimated and split into a donor (Cy3) and acceptor (Cy5) channel using a dichroic beamsplitter. Additional bandpass filters were placed before the detectors. Single-photon avalanche photodiodes (SPAD) (id100-50, low-noise, idQuantique, Switzerland) were used for detection. The output of the SPADs were inverted and directed into a time-correlated single-photon counting card (SPC150, Becker & Hickl GmBH). Samples were placed in 170 micron glass coverslip bottom 96-well microplates (Greiner Bio-One). A custom computer controlled microplate compatible x-y-z stage performs autofocusing and enables fully automated data collection. A Hamilton Microlab titrator (Microlab 500) automatically adds immersion water to the objective prior to each acquisition. Each FCS trace is the result of 10 × 300 s collections. FCS experiments were performed using 10 nM nucleosome bearing the FRET donor-acceptor pair in remodeling buffer (25mM HEPES, pH = 7.6, 0.2mM EDTA, 5mM MgCl2, 70mM KCl, 1mM DTT). The FCS measurements of the nucleosome in the presence SWR1C and the ATP analogue (AMP-PNP) were performed under saturating conditions. SWR1C was dialyzed overnight against the remodeling buffer prior to use in the FCS experiments. The acceptor autocorrelation function (G_AA_) and the donor/acceptor crosscorrelation function (G_DA_) were determined using Burst Analyzer software package (Becker & Hick1 GmBH). Since the relaxation time of the conformational fluctuation (observed rate constant, *k_obs_*) of the nucleosome can be derived from the ratio of any two correlation functions (Tims *et al.*, 2011; Torres and Levitus, 2007), we utilized the values of G_DA_/G_AA_ to obtain the kinetic parameters associated with conformational fluctuation of the nucleosome under various experimental conditions. The characteristic exponential curves associated with the ratio of two correlation functions (G_DA_/G_AA_) were analyzed using a single/double exponential rate equation, yielding the *k_obs_* values of the conformational fluctuation of the nucleosome.

### Transient kinetic measurements of nucleosome remodeling

The transient kinetic experiments of the SWR1C-catalyzed nucleosome remodeling reaction were carried out under single turnover conditions (excess SWR1C over nucleosome). The time-dependent fluorescence measurements during the SWR1C-catalyzed nucleosome reaction were carried out using an ISS PC1 spectrofluorometer. The nucleosome remodeling reaction were performed in remodeling buffer (25mM HEPES, pH = 7.6, 0.2mM EDTA, 5mM MgCl_2_, 70mM KCl, 1mM DTT) at 20 °C wherein the temperature was maintained using a circulating water bath. A representative nucleosome remodeling reaction contained 10nM nucleosome (bearing FRET pair), 60nm SWR1C, and 1mM ATP (or AMP-AMP). In order to monitor dimer exchange, a two-fold excess concentration of H2A.Z-H2B dimer (as compared to nucleosome) was used. The nucleosome was incubated with SWR1C in the presence or absence of the H2A.Z-H2B dimer for 5 min at 20°C to synchronize/pre-equilibrate the nucleosome-remodeler complex. The remodeling reaction was started with the addition of excess amount of ATP/AMP-PNP (1mM) which binds to the remodeler almost instantly (rapid-equilibrium). At least 3-4 kinetic traces were collected for each data set, and they were averaged to enhance the signal to noise ratio. The transient kinetic parameters of the SWR1C-catalyzed nucleosome reaction were obtained from the time-dependent change (loss/gain) in the Cy5 FRET signal (at 670nm upon excitation at 530nm). The averaged kinetic traces associated with the nucleosome remodeling reaction were analyzed using single or double exponential rate equations yielding the *k_obs_* values associated with the respective remodeling reaction. All curve fitting was performed in the Origin software package (OriginLab Corporation).

### ATPase Assays

The real-time and direct measurement of P_i_ was performed using a phosphate sensor (Brune *et al.* 1994). Our precise measurement of the pre-steady state kinetic parameters of SWR1C-catalyzed hydrolysis of ATP were unsuccessful even at the reduced temperature (4°C) and in the presence of lower concentration of ATP. The above experimental strategies, i.e., reduced temperature and lower ATP were, respectively, used to slow down the ATPase activity (for reliable rate measurements) and to reduce to the amount of free phosphate ion present in the ATP solution. In view of the above experimental limitation, we performed the steady-state kinetic analysis of the ATP hydrolysis catalyzed by SWR1C by discarding the initial data points of 300 sec. The experiments conditions used in the ATPase assay were as follows: SWR1C = 5nM, ATP = 100μM, H2A-nucleosome = 10nM, phosphate sensor = 2μM, H2A.Z-H2B dimer = 20nM. The real-time monitoring of p_i_ produced during the SWR1C-catalyzed reaction was performed on ISS PC1 spectrofluorometer upon exciting the sample at 425 nm and monitoring the emission as a function of time at 460nm. At least 3-4 kinetic traces were averaged and the resultant traces were analyzed using the steady-state equation as described below. (Fersht, 1999)

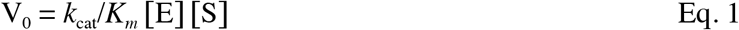

The amount of Pi produced during the steady-state of SWR1C-catalyzed ATP hydrolysis was calculated using the linear standard curve of Pi. The phosphate standard was purchased from Millipore.

---

**Figure S1.**
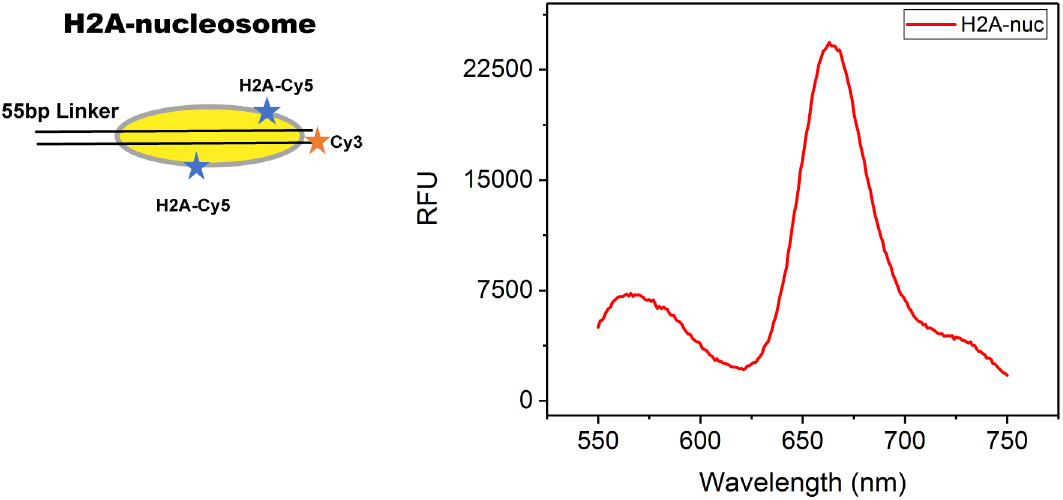
Emission spectrum of H2A-nucleosome containing Cy3-Cy5 as FRET pair. The emission spectrum (red trace) was collected using 10nM nucleosome in the remodeling buffer upon exciting the sample at 530nm.

**Figure S2.**
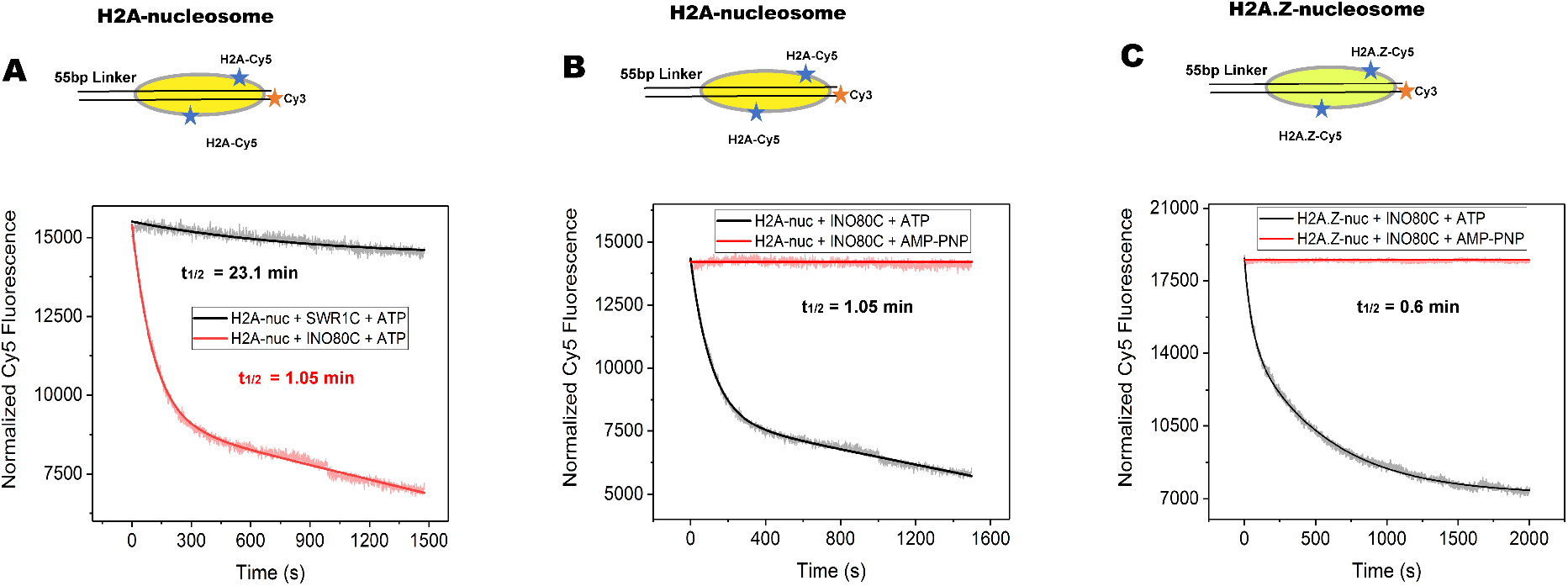
ATP-dependent nucleosomal DNA translocation catalyzed by SWR1C and INO80C. **(A)** ATP-dependent nucleosomal DNA translocation catalyzed by SWR1C (black trace) is markedly smaller than that of the INO80C (red trace), as evident from the relative amplitude of Cy5 FRET signal in the remodeling assay on the same substrate under identical experimental conditions. **(B) & (C)** INO80C catalyzes ATP-dependent translocation of the nucleosomal DNA on both the H2A-nucleosome and H2A.Z-nucleosome with an observed rate constant ~1.8 fold higher in the latter case.

**Figure S3.**
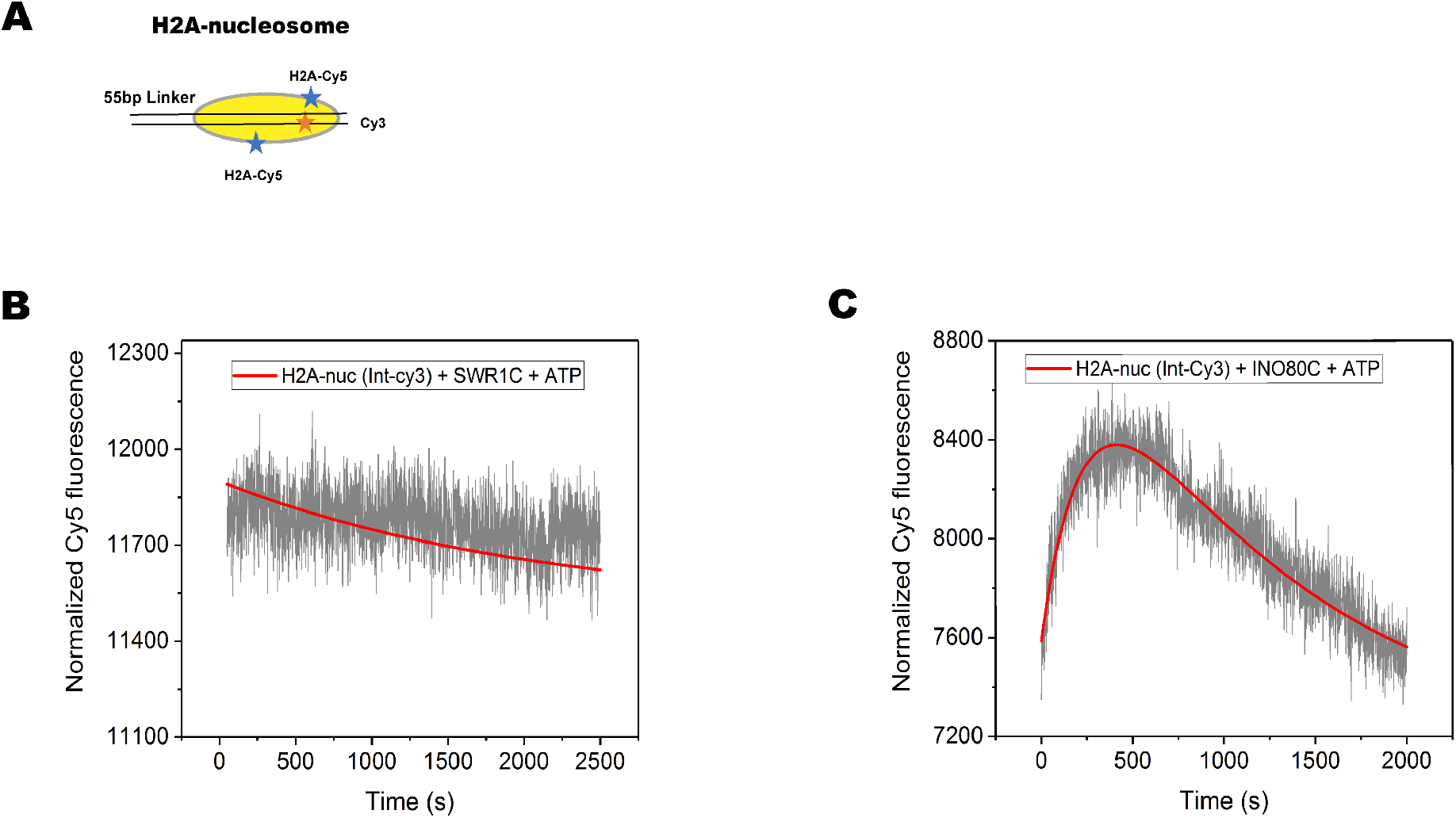
SWR1C catalyzes a slow translocation of DNA at the nucleosomal edge unlike INO80C which slides the end-positioned nucleosome towards center in a real-time. **(A)** Schematic of the nucleosomal substrate harboring FRET acceptor (Cy3) 15bp internal to the end of the nucleosomal DNA. **(B)** SWR1C catalyzes an ATP-dependent slow translocation of DNA on this nucleosome, as evident from a gradual loss of the Cy5 FRET signal. **(C)** INO80C, in contrast to SWR1C, catalyzes an ATP-dependent repositioning of the nucleosomal DNA leading to an initial enhancement in the Cy5 FRET signal followed by its gradual loss.

**Figure S4.**
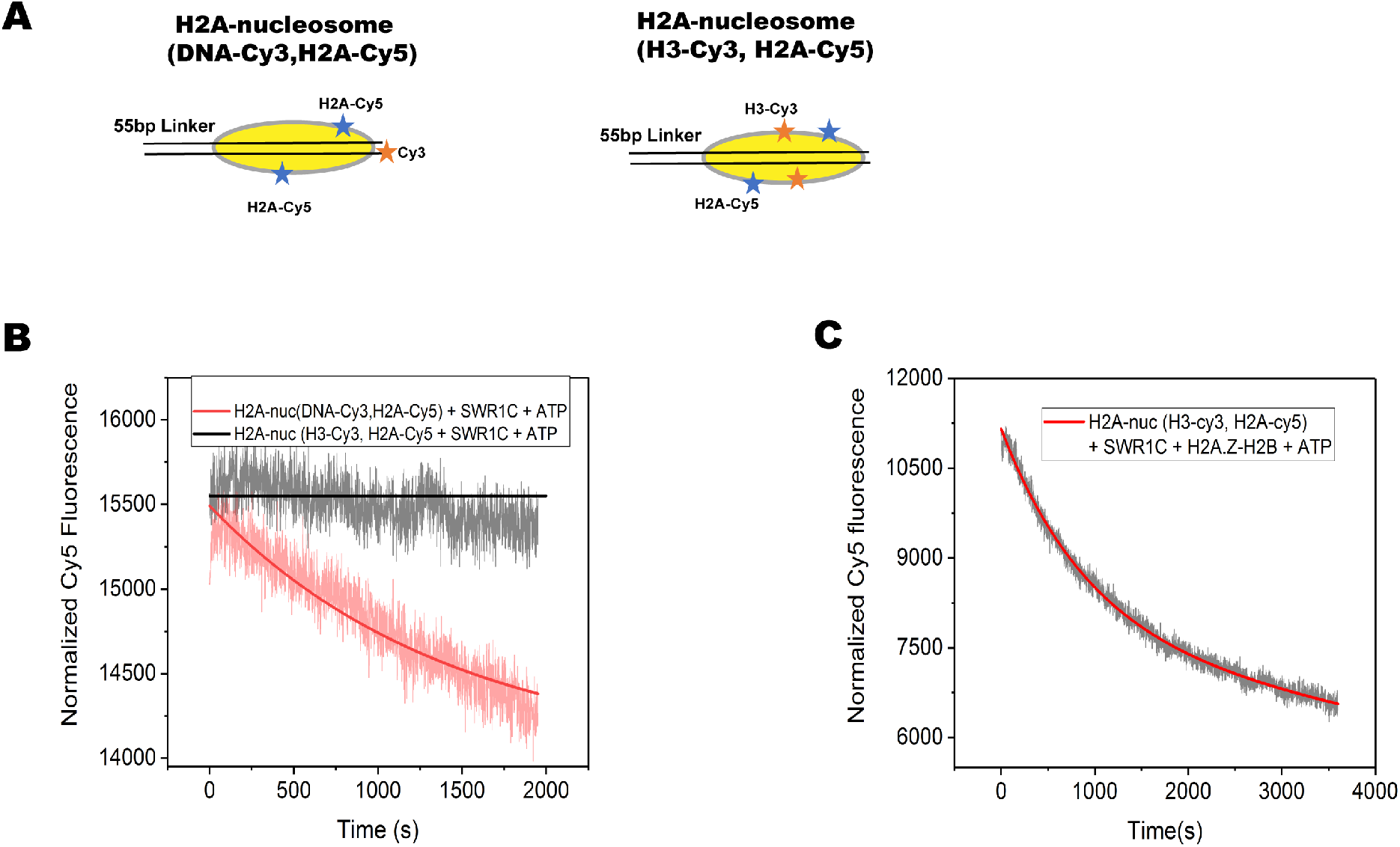
The SWR1C-catalyzed ATP-dependent translocation of nucleosomal DNA in the absence of the co-substrate (H2A.Z-H2B) does not facilitates the eviction of the H2A-H2B dimer. **(A)** Schematic of the nucleosomal substrates harboring Cy3-Cy5 FRET probe to monitor H2A-H2B eviction during DNA translocation and H2A-H2B eviction. **(B)** The ATP-dependent nucleosomal DNA translocation leads to loss of Cy5 FRET signal on the nucleosome bearing Cy3 on the nucleosomal DNA (red trace) as opposed to when it is located at H3-tail (black trace)—indicating that nucleosomal DNA translocation does <u>not</u> mediate H2A-H2B eviction from the H2A-nucleosome in the absence of H2A.Z-H2B dimer. **(C)** SWR1C catalyzes ATP-dependent eviction of H2A-H2B dimer from the H2A-nucleosome in the presence of its cosubstrate (H2A.Z-H2B).

**Figure S5.**
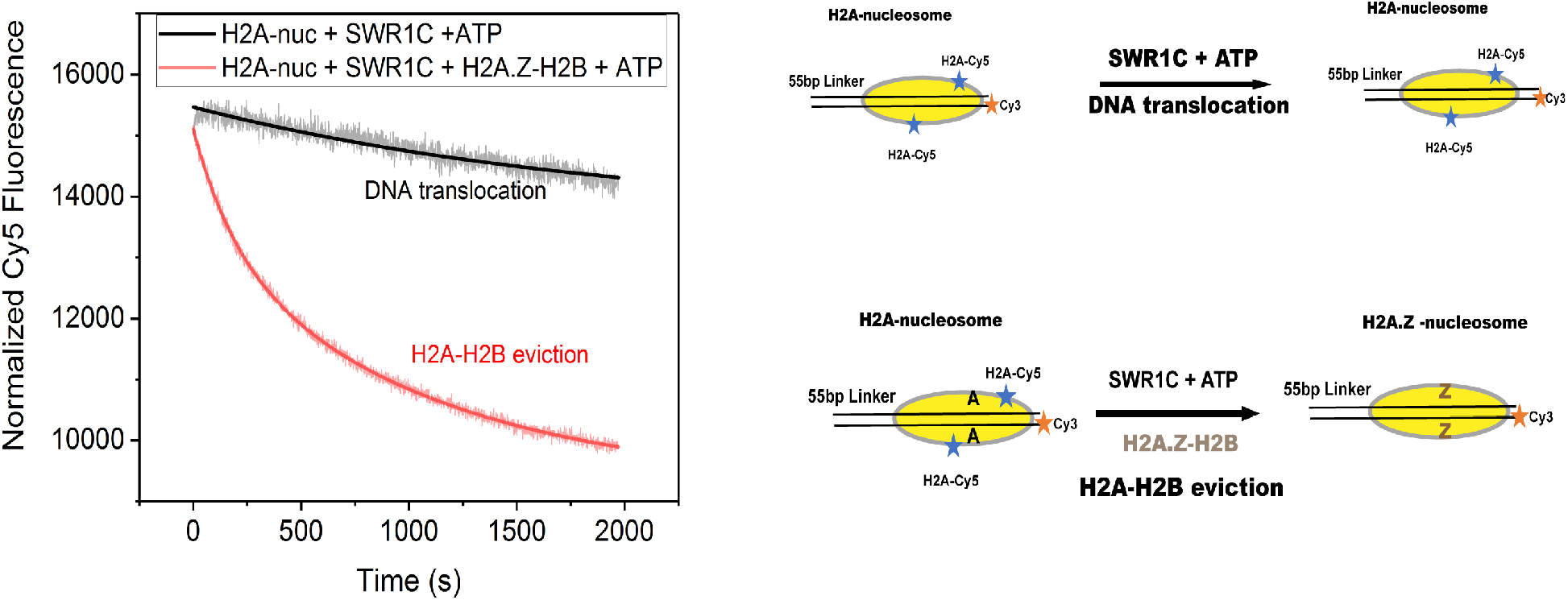
SWR1C catalyzes an ATP-dependent eviction of H2A-H2B from the H2A-nucleosome in the presence of its co-substrate (H2A.Z-H2B). In the absence of H2A.Z-H2B dimer, SWR1C catalyzes slow and localized translocation of the nucleosomal DNA (black trace), in contrast to a complete loss of the nucleosomal H2A-H2B dimer (red trace) – leading to a markedly greater change in Cy5 FRET signal in the latter case.

**Figure S6.**
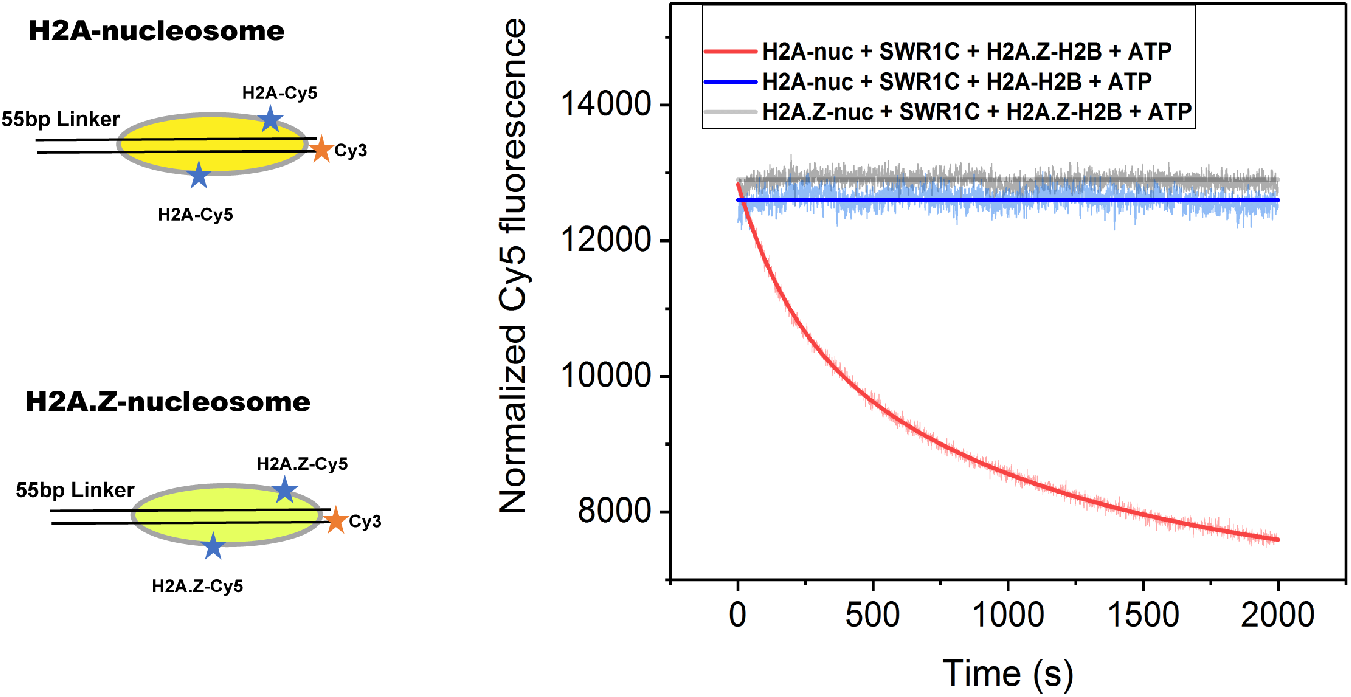
The substrate and the co-substrate discrimination by SWR1C in the ATP-dependent dimer-exchange reaction. The remodeler catalyzes the eviction of H2A-H2B from H2A-nuclesome only in the presence of H2A.Z-H2B as the co-substrate (red trace)—and not in the presence of H2A-H2B (blue trace). Additionally, SWR1C cannot catalyze an ATP-dependent eviction of H2A.Z-H2B dimer from H2A.Z-nucleosome.

